# A standardised framework to identify optimal animal models for efficacy assessment in drug development

**DOI:** 10.1101/382366

**Authors:** Guilherme S. Ferreira, Désirée Veening-Griffioen, Wouter Boon, Ellen Moors, Christine Gispen-de Wied, Huub Schellekens, Peter van Meer

## Abstract

**Introduction:** Poor translation of efficacy data derived from animal models is a potential contributor to costly and unnecessary attrition in clinical trials.

**Objectives:** To develop a tool to assess, validate and compare the clinical translatability of animal models used for the preliminary assessment of efficacy.

**Design and Results:** We conducted an exploratory literature search to identify the key aspects to validate animal models. Eight aspects (Epidemiology, Pathophysiology, Genetic, Biochemistry, Aetiology, Histology, Pharmacology and Endpoints) were identified for which questions were drafted to evaluate the different faces of the human disease simulation. Features of the framework include standardised instructions, a weighting and scoring system to compare models as well as contextualising factors regarding model similarity and evidence uncertainty. We included a quality assessment of the internal validity of drug intervention studies included in the Pharmacological validation section for both effective and ineffective drugs in humans. A web-based survey was conducted with experts from different stakeholders to gather input on the framework. Finally, we present a case study of a preliminary validation and comparison of two animal models for Duchenne Muscular Dystrophy (mdx mouse and GRMD dog) and Diabetes Type 2 (ZDF rat and db/db mouse). We show that there are significant differences between the mdx mouse and the GRMD dog, the latter mimicking the human condition to a greater extent than the mouse despite the considerable lack of published data. In DT2, both the ZDF rat and the db/db mouse are comparable with minor differences in pathophysiology.

**Conclusions:** FIMD facilitates drug development by serving as the basis to select the most relevant model that can provide meaningful and translatable results to progress drug candidates to the clinic.

## Introduction

The use of animals to evaluate the safety of new drugs is an integral part of the regulatory research and development process (1,2). Established at a time when laboratory animals were one of the most complex systems available, they are still considered as the gold standard today. Yet, despite their apparent value as a drug testing system to predict safety and efficacy in humans, scientists are increasingly aware of their considerable drawbacks and limited predictivity (3–6).

While no apparent toxicity in poorly predictive animal models can lead to possible harm to patients, false toxic signals might prevent potentially safe drugs from reaching patients. This constitutes an unreasonable loss of resources for drug developers. Concomitantly, limitations in animal models of efficacy showing an overly optimistic interpretation of efficacy will lead to clinical trials with drugs that have a modest effect at best or are completely ineffective at worst (5,7).

We previously assessed the value of regulatory safety studies in a public-private research consortium which consisted of pharmaceutical company stakeholders, the Dutch regulatory agency and academia (8). This partnership was unique in that it allowed proprietary data to be used for our primary analyses, which could then be presented in an aggregated, anonymised fashion to propose policy changes (9–13). A key finding of these studies was that despite non-human primate (NPH) models having the closest biological resemblance to humans, their indiscriminate use in the safety testing of new biotechnology products (e.g. mAbs) as well as in demonstrating the similarity of biosimilar to reference products often adds limited value to the preclinical package. When taken altogether, these results suggest the mandatory use of animal safety testing according to current guidelines should be reconsidered.

Contrary to safety assessment, the evaluation of efficacy is not subject to formalized guidance or regulations since each new drug warrants a tailor-made approach based on its mechanism of action and indication (14). Consequently, predefining which assays or models to be used to test new drugs’ efficacy, as done for safety, could jeopardise innovative companies’ ability to develop such drug-specific strategies.

Nevertheless, most late-stage clinical trials, which are often based on efficacy data from animal studies fail due to the lack of efficacy (15–19). The low internal validity (i.e. the methodological qualities of an experiment, such as randomisation and blinding) of animal research has been frequently suggested as a likely cause for such poor translation to the clinic (20–24). Initiatives aimed at improving design and reporting standards of preclinical studies, such as the ARRIVE guidelines, now allow researchers to effectively address these issues (25).

The inadequate assessment of the external validity of efficacy models (i.e. how well animal results are generalisable to the human situation) is also an important factor for poor translation (4). Currently, drug developers frequently rely on the well-established criteria of face, construct and predictive validity (26,27). Because none of these criteria goes beyond the level of scientific concept – they are not integrated and do not present a systematic way to assess the ability of an animal model to predict drug efficacy in humans, they are highly subject to user interpretation. The absence of standardisation results in animal models being assessed by different disease parameters, which further complicates a scientifically relevant comparison.

Previous attempts to formalize the assessment of external validity have introduced systematic ways to score validity, but fail to capture most of the characteristics potentially relevant for the demonstration of efficacy (e.g. genes, biomarkers, histology) to make them informative and usable to this end (28,29).

### The Framework to Identify Models of Disease (FIMD)

The existing approaches for assessing external validity cannot be used by researchers to find what is the most relevant model to demonstrate the efficacy of a drug based on its mechanism of action and/or indication. Here, we present a method to assess the external validity of efficacy models as well as to integrate the different aspects needed to establish preliminary efficacy in an animal model – the Framework to Identify Models of Disease (FIMD). A ‘model of disease’ is here used for any animal model that simulates a human condition for which a drug can be developed. Eight relevant aspects were identified based on an exploratory literature search (see Supplementary Information S1) and used to draft questions related to the different facets of disease simulation (Fig 1).

**Fig 1.**
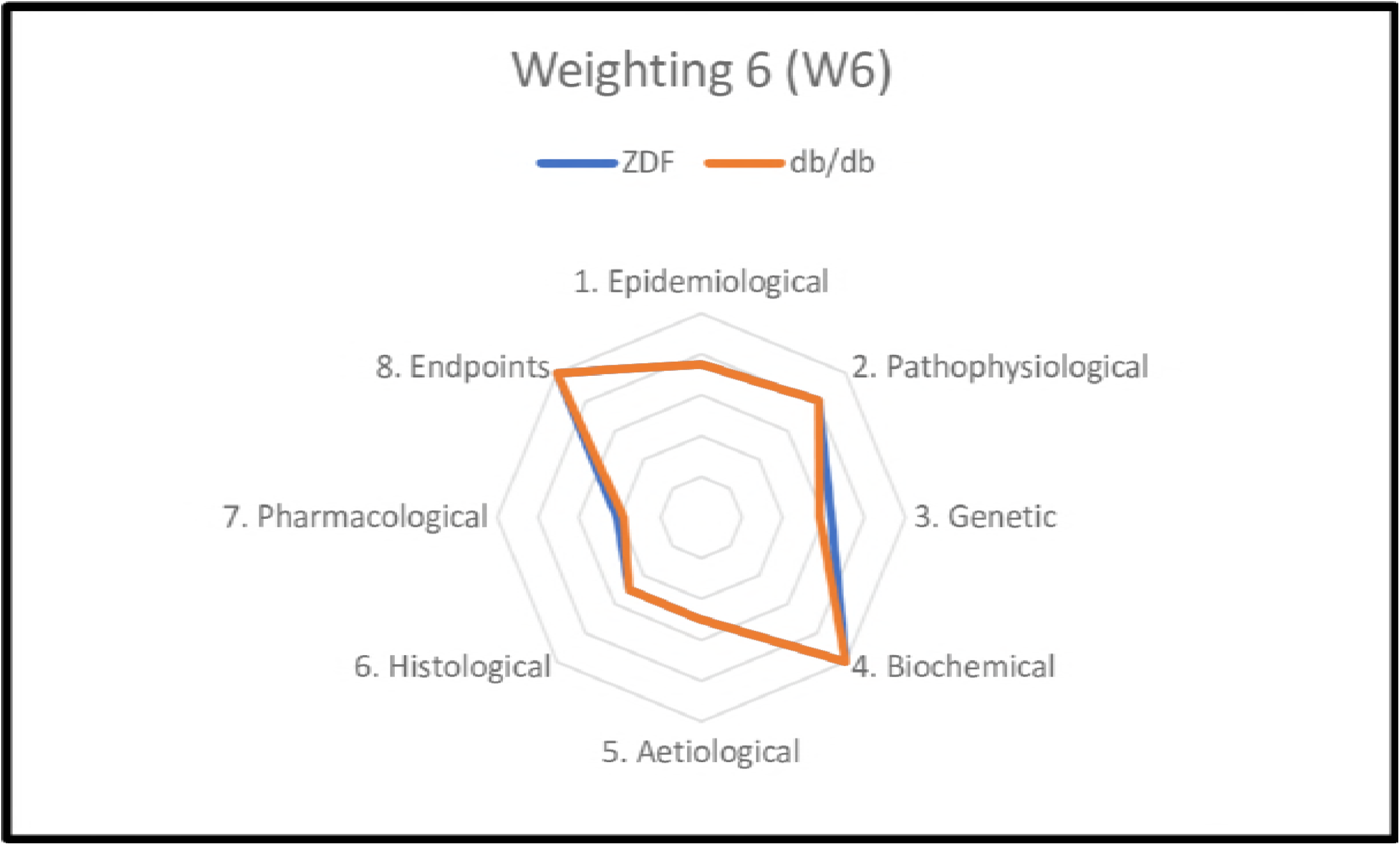
The Framework to Identify Models of Disease (FIMD). Questions per validation parameter.

We designed FIMD to circumvent some important limitations present in the current approaches and further identified in a web-based survey with experts from academia, industry and regulatory agencies (see Supplementary Information S2). To make it systematic, transparent and to minimise parameter discordance, instructions on how to complete the validation sheets are provided (see Supplementary Information S2). Animal studies of both effective and ineffective drugs are included. This results in a better understanding of the pathways that are involved in the disease pathophysiology of a model when compared to humans. Consequently, companies with extensive animal data from failed projects can easily perform a read-across of their models to inform the choice of future programmes. In addition, all interventional drug studies in the pharmacological validation also include a quality assessment of the study design and reporting adapted from the ARRIVE guidelines (see Supplementary Information S2).

To facilitate the comparison between animal models in the same indication and ultimately, the choice of the best fit for investigating a drug’s efficacy, we developed a weighting and scoring system (see Supplementary Information S3). This system includes a Disease Classification Flowchart (DCF) for the selection of the adequate weighting system since the relevance of each aspect might differ for different indications (e.g. genetic disorder vs. bacterial infection).

The weighting and scoring system is an indication of the degree to which a model simulates the human condition. While it is admittedly arbitrary, it allows researchers to identify the strengths and weaknesses of an animal model at a glance. The underlying data will further determine which model is the most relevant for a given drug. This means the best fit will not necessarily be the model with the highest score but rather the model that more closely mimics the pathways involved in the mechanism of action of a drug.

Next to the weighting and scoring system, two factors were created to further contextualise the final score: the uncertainty factor and the similarity factor. The uncertainty factor differentiates between models that are not well-characterised and models that scored low for not simulating completely or partially many aspects of the human condition. The similarity factor differentiates between two models with similar final scores but that score differently in the same aspects of validation.

All these features of the framework were further refined the framework, we conducted a web-based survey with experts from academia, industry and regulatory agencies (see Supplementary Information S4).

### Validating Efficacy Models in FIMD

We also addressed the need to clearly define what are the minimum requirements to validate an animal model. There are several interpretations of what the validation of an assay or model should entail and how to assure its reproducibility. For the OECD and EURL-ECVAM, validation means establishing a statistically rigid range for several criteria to describe a test’s ability to be reproduced reliably (30). This is often an expensive and lengthy process, mostly applicable to *in vitro* tests, which would not be practical or add much value to efficacy assessment in animal models (14). A more applied approach is presented by ICHS5(R3), in which the definition of the context of use (i.e. the conditions in which the assay results can be relied upon) is the guiding factor for the validation of a test (31).

Our validation definition aims to provide the evidence for an efficacy model’s context of use (i.e. the intended indication for which drugs are being developed). Although animal models are not expected to completely mimic the human condition, it is important to identify which aspects they can reproduce and to which extent. Therefore, we established four levels of confidence in the validation of animal models in their context of use based on the percentage of definite answers to the eight validation sections (see Table 1). A ‘definite answer’ is defined as any answer except for ‘unclear’, which is used to indicate the absence of evidence in the literature or conflicting results.

**Table 1:**
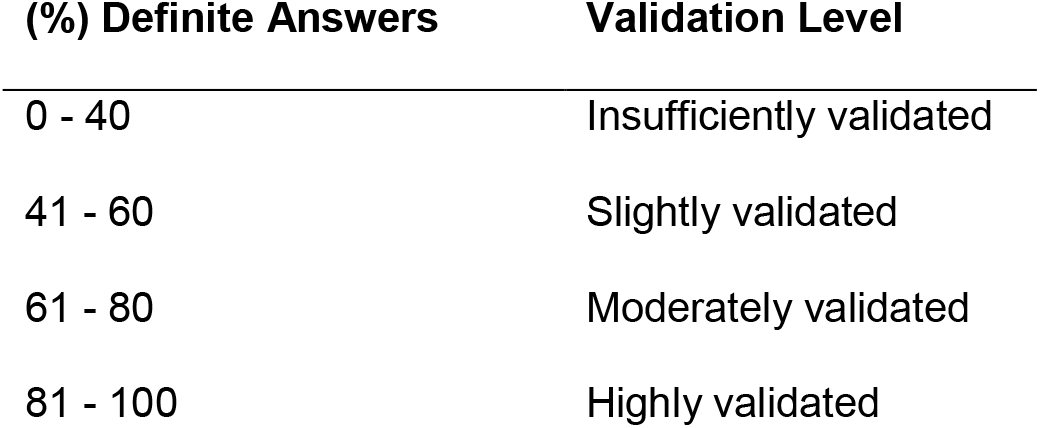
Different levels of validation according to the percentage of definite answers to the questionnaire.

### Applications and Final Considerations

The product of FIMD is a validation sheet of an animal model for an indication, which provides the necessary information for its assessment as a potential model to demonstrate a drug’s efficacy (see Supplementary Information S4, S5 and S6). Based on a drug’s mechanism of action, models can be first discriminated by assessing whether the correlation between animal and human drug studies of relevant pathways is available in the pharmacological validation section. The other sections provide additional information, such as the presence of relevant genes and biomarkers. Finally, the validation level is an index of the reliability of a model’s overall ability to mimic the human condition, serving as another layer to further differentiate potentially useful from non-useful models. The combination of all these features allows researchers to select, among a plethora of models, the model most likely to correctly predict the efficacy of a drug in humans. An example of the application of FIMD is presented in Box 1, in which the Golden Retriever Muscular Dystrophy (GRMD) dog emerges as a significantly better model than the more commonly used mdx mouse.

An important application for FIMD is on the approval of animal studies by Institutional Review Boards (IRBs). IRBs often base their decisions on unpublished animal studies with poor internal validity (32). FIMD presents an opportunity to assess the validity of a specific model for efficacy assessment while also providing insights from earlier research. By using the validation sheet of models used to support the first-in-human trials, IRBs can, for the first time, tackle all these issues at once. FIMD provides the background for the choice of the model(s), allowing IRBs to accurately assess whether the data generated is likely to be translatable to the clinic.

Since FIMD includes a quality assessment of all studies included in the pharmacological validated and require that these be published, it promotes further scientific scrutiny in peer-review processes. Nonetheless, it only includes publicly available information. Given the publication bias often reported in animal and clinical research, it is possible that results from the pharmacological validation might be skewed (4,33,34). Nonetheless, there is a growing demand for the pre-registration of preclinical studies and the publication of their results (32,35,36). With the establishment of registries like PreclinicalTrials.eu, the overall publication bias is expected to be reduced and therefore, so will be its impact on FIMD (37). Furthermore, with the collection of validation sheets of models of efficacy, it will be possible to make them available in an open database of validated models in which users can, based on their drug’s characteristics (e.g. mechanism of action or intended indication), find the best model to evaluate the efficacy of a drug before planning an animal experiment.

FIMD can simplify the interaction between companies and regulatory agencies as it allows a more objective and science-based discussion on the choice of an animal model. This potentially prevents efficacy studies on non-relevant models, effectively contributing to the reduction of the use of animal models in the context of the 3R’s. By assessing the degree to which a model mimics a human condition, FIMD facilitates the choice of a relevant model for efficacy assessment and promotes the conduct of efficacy studies whose results will more likely translate to the clinical situation.

##### Box 1: Using FIMD to assess and compare efficacy models

We used a simplified version of FIMD to test its applicability in two indications with two models each (more information available in Supplementary Information S4). Duchenne Muscular Dystrophy (DMD) was chosen because it is caused by mutations in a single gene and for the limited availability of effective therapies (38). Type 2 Diabetes (T2D) was chosen as a disease which has a complex pathophysiology and for which an extensive set of therapies is available (39).

In DMD, the models were chosen based on being either commonly used (mdx mouse) or for being perceived as offering a better replication of symptoms and histological features (Golden Retriever Muscular Dystrophy – GRMD – dog) (40,41). A total of 58 articles were included for the mdx mouse and 41 for the GRMD dog. The relative scores per parameter are presented in Fig 2.

**Fig 2.**
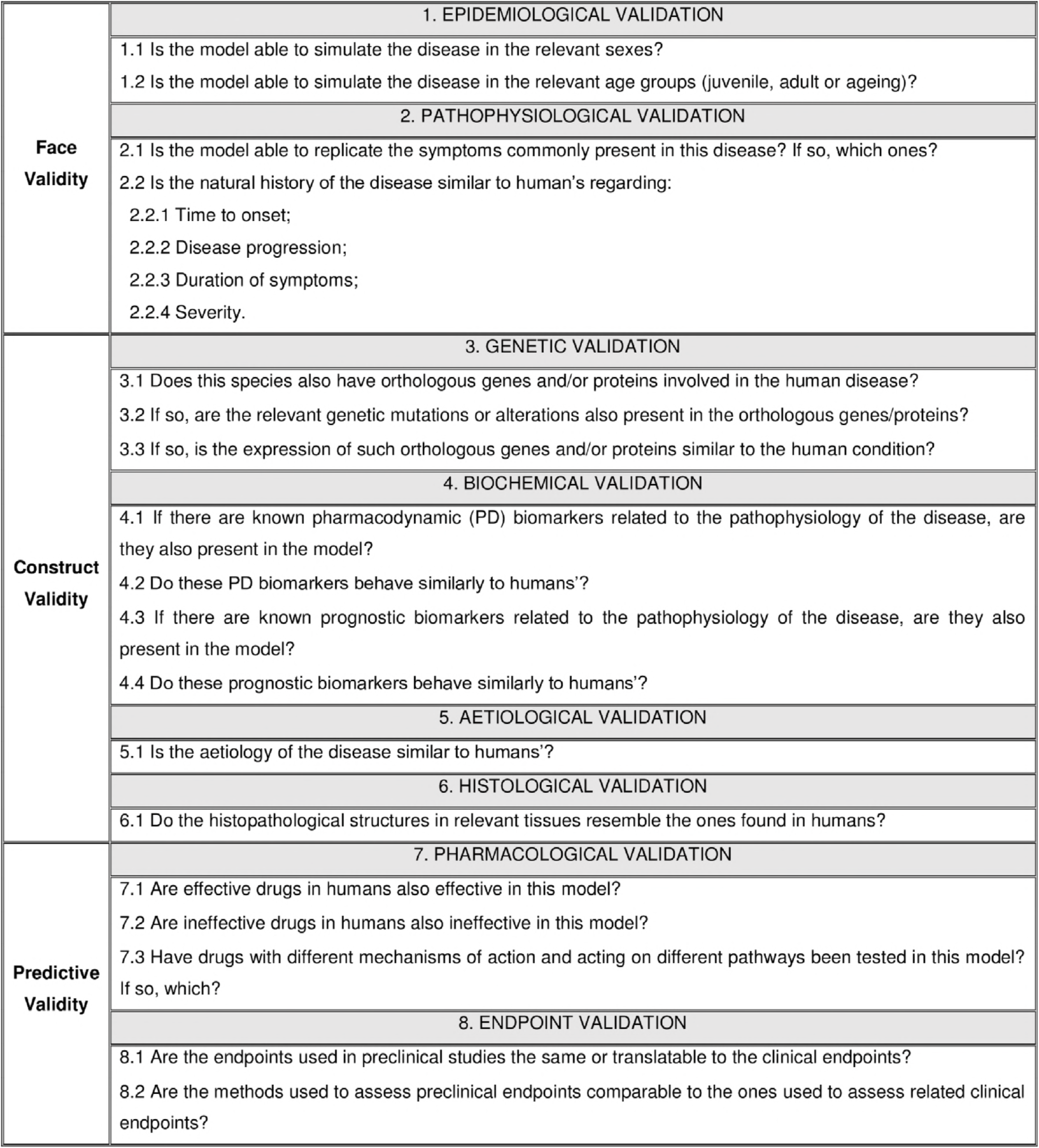
DMD models results. Radar graph showing the scores per parameter per model of DMD for weighting 2. The closer a parameter is to the edge, the better the model simulates that specific aspect of the human disease.

The GRMD dog scores better in the epidemiological, pathophysiological and histological sections while the mdx mouse does so in the pharmacological and endpoints sections. The GRMD dog mimics the natural history of the disease, symptoms (e.g. muscle wasting) and histopathological features (e.g. muscle regeneration) better than the mdx mouse. Especially for drugs which aim to slow down muscle degeneration or delay the disease onset, the GRMD dog is likely to generate more translatable data than the mdx mouse.

The difference in the pharmacological and endpoints sections stems mostly from the uncertainty factor, which is 20% and 5.7% for the GRMD dog and the mdx mouse respectively. Since most drug screening studies are done in mdx mice, there are more studies available for the pharmacological validation. However, there are no published studies in GRMD dogs for most drugs tested in humans. The only published study that assessed a functional outcome, did so in sedated dogs, which reduced the score of the endpoints validation. Hence, a comparison between these models in these two sections is unlikely to be informative.

For T2D, the Zucker Diabetic Fatty (ZDF) rat and the db/db mouse were both chosen for being routinely used in drug screening for antidiabetic drugs. A total of 195 publications were included for the ZDF rat and 282 for the db/db mouse. The relative scores per parameter are presented in Fig 3.

**Fig 3.**
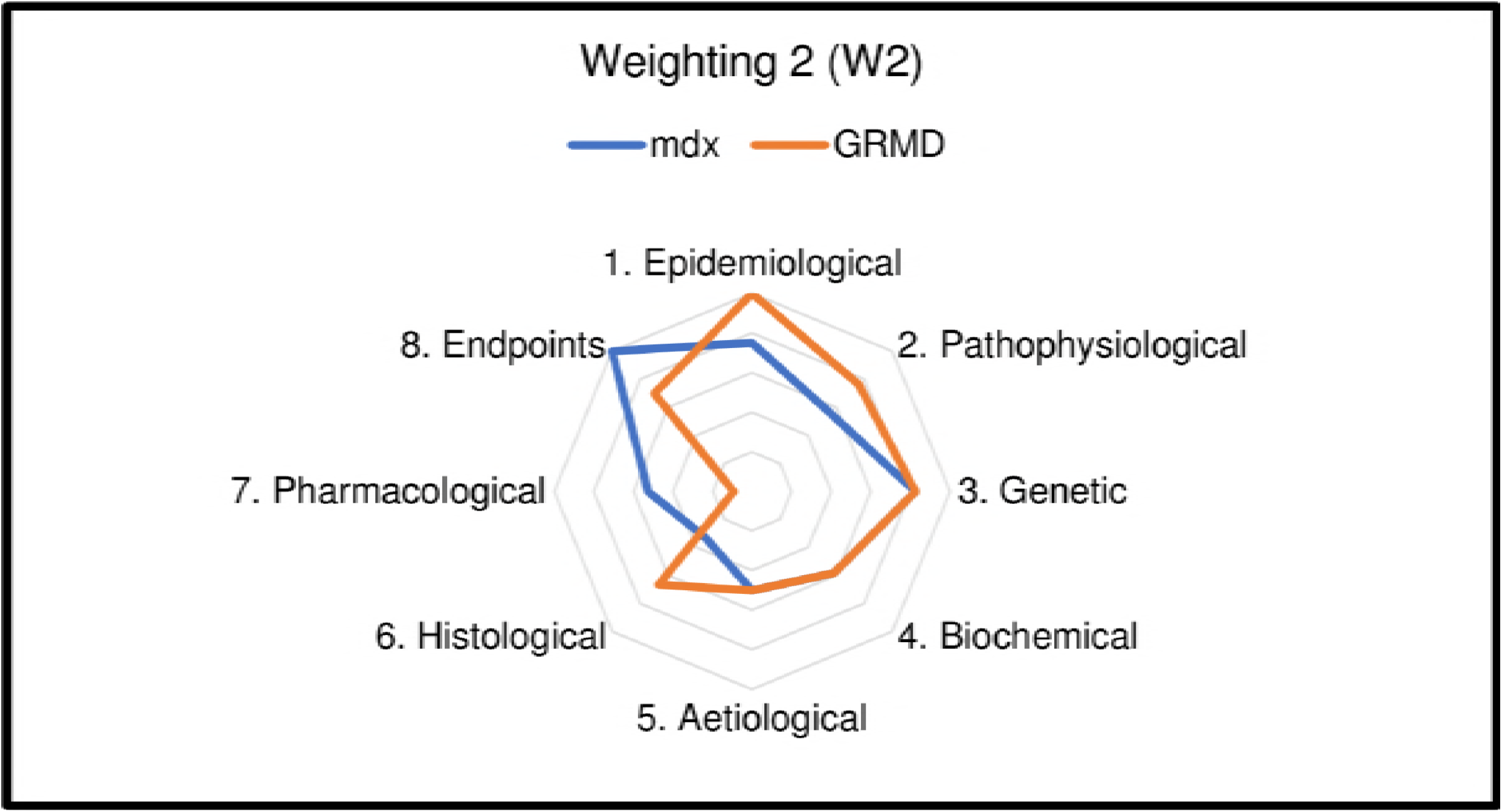
T2D models results. Radar graph showing the scores per parameter per model of T2D for weighting 6. The closer a parameter is to the edge, the better the model simulates that specific aspect of the human disease.

Compared to the DMD models, there are only minor differences since the similarity factor between the two models is almost 90%. Both T2D models have uncertainty factors of over 20% due to the lack of studies on genetic characterisation and in the pharmacological validation. [/BOX 1]

## Acknowledgements

We thank all the participants of the web-based expert panel (which was anonymous) and Dr Elaine Teixeira Rabello for her constructive comments on the manuscript.

